# Dynamic tracking of variant frequencies depicts the evolution of mutation sites amongst SARS-CoV-2 genomes from India

**DOI:** 10.1101/2020.07.14.201905

**Authors:** Shenu Hudson B., Vaishnavi Kolte, Azra Khan, Gaurav Sharma

## Abstract

With the exponential spread of the COVID-19 pandemic across the world within the twelve months, SARS-CoV-2 strains are continuously trying to adapt themselves in the host environment by random mutations. While doing so, some variants with evolutionary advantages such as better human to human transmissibility potential might get naturally selected. This short communication demonstrates how the mutation frequency patterns are evolving in 2,457 SAR-CoV-2 strains isolated from COVID-19 patients across diverse Indian states. We have identified 19 such variants showing contrasting mutational probabilities in the span of seven months. Out of these, 14 variants are showing increasing mutational probabilities suggesting their propagation with time due to their unexplored evolutionary advantages. Whereas mutational probabilities of five variants have significantly decreased in June onwards as compared to March/April, suggesting their termination with time. Further in-depth investigation of these identified variants will provide valuable knowledge about the evolution, infection strategies, transmission rates, and epidemiology of SARS-CoV-2.

## Introduction

Since the emergence of SARS-CoV-2 in Wuhan, China in December 2019, the infection has spread at a menacing rate throughout the world. As of today, over 70 million active cases of COVID-19 and 1.5 million deaths are telling the horror story of this debilitating virus. Since the whole genome sequencing of the SARS-CoV-2 Wuhan Hu-1 strain earlier this year, more than 250,000 genome sequences ^1–3^ have been deposited in open-source platforms such as GISAID, NCBI Virus, etc. Consortiums such as Nextstrain are providing valuable, unprecedented insights into the demography and epidemiology of SARS-CoV-2 strains along with information to supervise drug/vaccine design and discovery.

Coronaviruses encode a 3′-5′ exoribonuclease (Nsp14) that proofreads RNA ^4^, which tries to maintain genome fidelity and control variations, although even with this, mutation rates amongst these viruses are very high. However, during the past twelve months of the ongoing COVID-19 pandemic, worldwide human-to-human transmission has enabled SARS-CoV-2 to accumulate numerous genetic variations ^5^. Several mutations among these are non-significant while other mutations could be beneficial for the survival of the virus, its infecting capability, and transmission ^6–9^. Given that beneficial mutations could be naturally selected in wider populations, studying SARS-CoV-2 genomic variants and their tracking with time might help us in understanding viral evolution, behavior, and infection trajectory.

In this study, we have tracked and identified several mutational sites for month-wise (March to September; seven months) separated SARS-CoV-2 genome datasets isolated from Indian COVID-19 patients to distinguish which variants are getting naturally selected to propagate further and the ones which are being dismissed from the population.

## Methods

2,457 complete genomes of SARS-CoV-2 isolated from Indian patients till September 30, 2020, were extracted from GISAID on November 1, 2020 **(Table S1)**. From September 30 to date, only eight Indian SARS-CoV-2 genome sequences have been submitted to GISAID, owing to this, we have not used genome data after September month in this study. Based on the sample collection date of each strain, we distributed them into seven categories: ‘March’ (this included the strains collected in January, February, and March), April, May, June, July, August, and September. The genome of the hCoV-19/Wuhan/WH01/2019 strain (EPI_ISL_406798) was used as a reference throughout this analysis ^3^. For each category, we generated multiple sequence alignment and identified the percentage of all four nucleotides (along with gap (-) and other non-standard nucleotides) at each site. Sites having >2% gaps or non-standard nucleotides were not considered in the analysis. The frequency of each nucleotide in the alignment was calculated and a ratio was determined with the frequency of the nucleotide in the reference genome. Mutation frequency/probability was defined as the ratio of the frequency of the nucleotide at any site to the frequency of the nucleotide present in the reference sequence at the same aligned site. A hierarchical clustering based heatmap of each nucleotide loci was generated using mutational probabilities within each category using the hclust function in R. Simultaneously, trend plots were also generated for all identified clusters using ggplot.

## Results

All 2,457 SARS-CoV-2 strains isolated from the diverse landscape of India during March-September 2020 were categorized into month categories (**Table S1)** and aligned with the reference WH01 genome. This study recognized 268 sites with mutation probability ranging from 2 to 97.99% in at least one month. We identified that in most of these sites, there were negligible variations (>2% and <4%) amongst different month categories. Therefore, to identify the critical variations in our downstream analysis, we used a 4% minimum mutational probability score in at least one-month category and found 118 sites encompassing this criterion. Accordingly, we identified 36, 33, 32, 36, 37, 37, and 50 highly mutating sites amongst the March, April, May, June, July, August, September categories, respectively. Of these, 11 sites were showing significant mutation probabilities in all seven months, 3 in six of the months, 8 in four of the months, 11 in three categories, 16 in two, and 69 were unique in just a single month category. Finally, the mutational probabilities of these 118 sites across all time points (month-wise) were visualized using a clustered heatmap where six clusters were obtained with 4, 3, 2, 5, 5, and 99 sites per cluster **(Figure 1AB, Table S2)**. The largest cluster with 99 sites did not show an upward or downward longitudinal trend, therefore, it might be classified as a “neutral” cluster **(Figure 1AB)**, whereas 19 sites distributed in other five clusters showed significant variations, which are discussed below.

**Figure 1:**
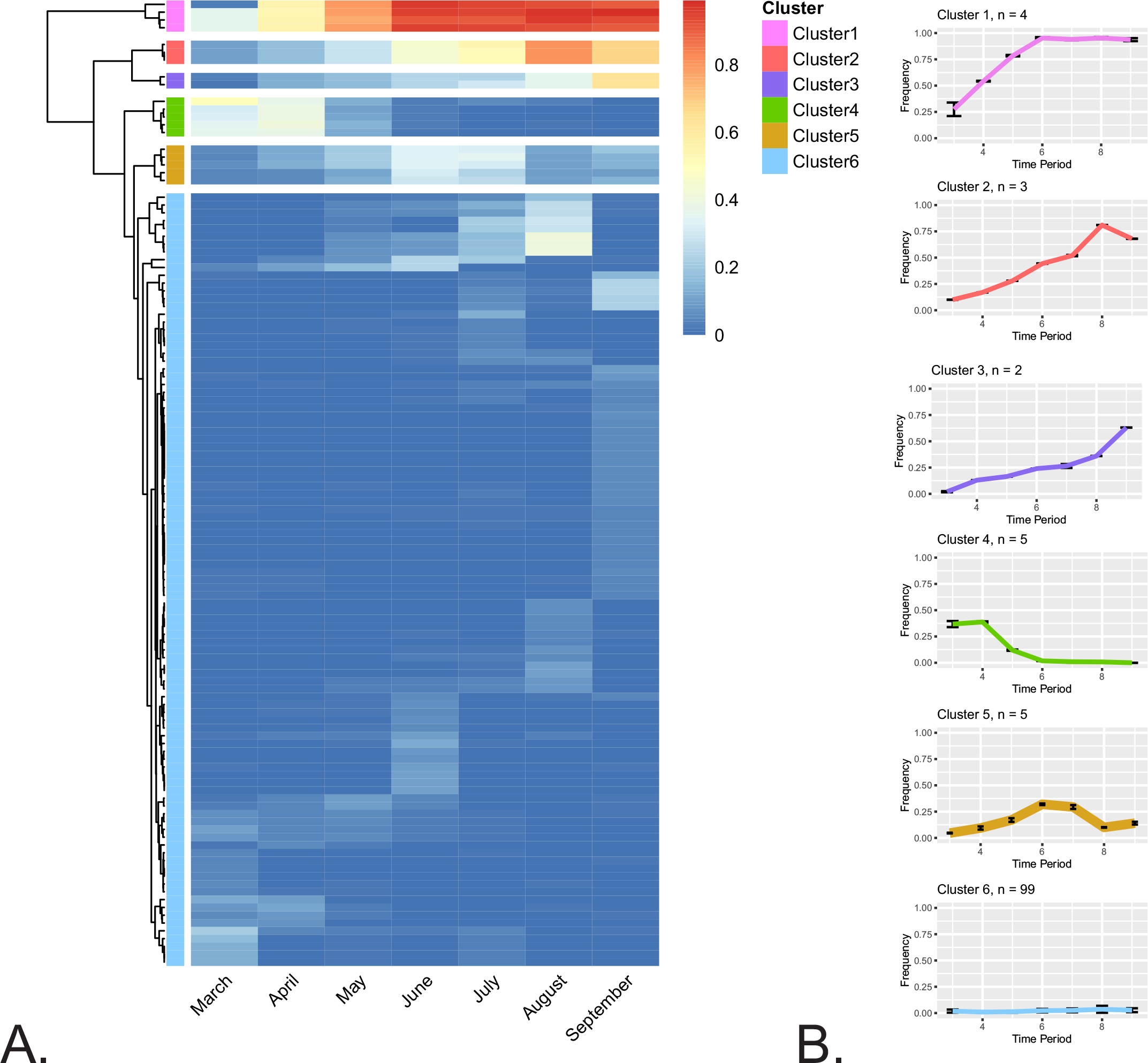
Dynamic tracking of mutational frequencies within genomes extracted from Indian samples in March, April, May, June, July, August, and September. **A) Heatmap of mutational probabilities. B) Trend plots of average mutational probabilities within each cluster**. For each cluster, n represents the number of sites. Time period on X-axis denotes month categories.

**Cluster one** included four loci where the mutational probabilities were lowest in March (2-35%) with a radical increase to >90% in June and onwards, indicative of positive selection in the population (**Figure 1AB, Table S2**). The first site, C3037T is in the nsp3 protein-coding region of the orf1a gene, C14408T is in RdRp (RNA-dependent RNA polymerase) protein-coding region within the orf1ab gene, the third site (A23403G) is in the spike protein-coding gene, and the fourth site (C241T) is in the noncoding 5’ UTR region. C3037T is a synonymous mutation (F106F in nsp3 protein) and does not lead to any major changes in protein structure and function. On the other hand, we noticed that C14408T mutation causes a nonsynonymous change (corresponding to P125L in RdRp) which has also been observed in genomes isolated from different continents suggested to change the rigidify of RdRp structure ^10^. The third site, A23403G, is well known to cause a D614G mutation in the spike protein, which interferes in domain S1-S2 interaction. This has been suggested to cause substantial conformational shifts in spike protein that in turn enhance SARS-CoV-2 infectivity while retaining sensitivity to antibodies that target the receptor-binding domain ^6,7^. Sites identified in the **second cluster** started showing mutations in April and in each month their frequency is expanding, reaching up to 60-80% of the total population in August/September. This cluster consists of three sites, of which one is synonymous mutations **(Table S2)** while the other two are non-synonymous and worth exploring further. All three sites G28881A, G28882A, and G28883C are located consecutively in the nucleocapsid gene and lead to changes in amino acids R203K, R203R, and G204R, respectively.

**Cluster three** shows a trend like cluster two however the increase in percentage representation is slow and goes up to 35-60% as compared to 60-80% in the latter. It consists of two sites only i.e., a synonymous mutation in C313T [nsp1 in ORF1a; L16L] and a non-synonymous mutation in C5700A [nsp3 in ORF1a; A994D]. The **fourth cluster** consists of five sites that had high mutational probabilities during early infection months, *i*.*e*., March and April, and decreased mutational probabilities in May and June, and further decreasing to zero in August and September, indicating that these variants are not being selected for further propagation. Two out of these sites, C13730T [which results in an A4489V change in ORF1ab (A97V in RdRp protein)] and C23929T [which results in a Y789Y change in spike protein], either maintain the general chemical nature of the amino acid (Alanine to Valine change) or are fully synonymous. However, the other three sites, C6312A, G11083T, and C28311T, result in non-synonymous changes in the nsp3 region (T1198K) of the ORF1a (T2016K), the nsp6 region (L37F) of ORF1a (L3606F), and the nucleocapsid (P13L) protein, respectively. Based on trends, we can hypothesize that the later three non-synonymous mutations were not selected by the virus population for further propagation on account of their putative lesser efficiency in infection or other types of fitness disadvantages.

The **fifth cluster** consists of five sites where the mutational probabilities were making a bell-shaped curve across all month-wise datasets meaning that in March and April along with August and September, the mutational frequencies were in the range of 2-10%, however, we could see >30% mutational representation in June and July months. G25563T site is a part of the orf3a gene and leads to Q57H non-synonymous mutation. Similarly, another non-synonymous mutation was identified in C28854T that leads to S194L variation in Nucleocapsid protein. Other loci include synonymous mutations at C18877T [nsp14], C22444T [S], and C26735T [M], which cause insignificant variations as indicated by L280L, D294D, and Y71Y, respectively. Most of the **sixth cluster** sites have small variations ranging from 0-10% mutational representations in one of the months. However, in a few sporadic sites, the representation was 10-40% in only one of the months and in other categories, the representation was non-significant suggesting the absence of any pattern. Finally, considering the ongoing trends of these variants, we hypothesize that sites identified in clusters 1, 2, and 3 must be getting selected for propagation owing to some unknown fitness advantages. Similarly, as the representation of sites identified in clusters 4 and 5 was higher earlier and at its lowest in the end months, we argue that these sites are not getting preferred during viral population selection.

To counter evolutionary pressure, viruses, akin to other living beings, continuously mutate their genetic material to improve their infection strategies, resistance potential to antiviral therapies, and transmission rate ^11–13^. Most of these random mutations are synonymous or functionally insignificant. However, a few non-synonymous mutations might give an extra advantage to the virus in its faster transmission, additional infection severity, or higher resistance against antiviral vaccines/treatments along with other fitness advantages. Overall, this study suggests that all identified mutations are not evenly distributed across the virus population during different timeframes; however, some loci are more prone to propagate and some get terminated with time. As several variants in clusters 1, 2, and 3 have higher mutational probability in August/September as compared to March/April, understanding the consequences of those propagating variants in terms of infection and epidemiology will be of great importance. Exploring and recognizing this information might prove helpful for drug and vaccine development. Some reports have already shown the rapid increase of a non-synonymous (D614G) variant across the globe that might have facilitated increased human-to-human transmission ^6,7^. In-depth investigations of variants identified in this study will provide newer insights into the evolution and fitness advantages acquired by SARS-CoV-2.

## Declarations

### Ethics approval and consent to participate

Not applicable

## Consent for publication

Not applicable.

## Availability of data and materials

All data and materials are included in this published article.

## Authors’ contributions

GS conceived and designed the study and wrote the paper. SHB, VK, AK, and GS performed the study and analyzed the data. All authors read and approved the final manuscript.

## Acknowledgments

GS would like to thank the Department of Science and Technology (DST) and IBAB, Bengaluru for financial and infrastructure support. The authors would like to thank his colleague Dr. Shruthi S. Vembar for constructive criticism of the manuscript. The authors would also like to thank Dr. R. Srivatsan for his help in statistical analysis. We would also like to thank the GISAID database for allowing us to access the genome sequences for this scientific research.

## Legends for Supplementary Tables

**Table S1:** Location-based distribution of 2,457 Indian SARS-CoV-2 strains analyzed in this study.

**Table S2:** Mutation probability scores and other relevant metadata for 268 identified mutation sites at Genome, Codon, and protein level. Heatmap-based cluster information using 4% minimum mutational probability data (118 sites) is also provided in the table. NA represents “Not Applicable” for cluster information as those sites were not considered for heat map-based clustering.

